# PhyMapNet: A Phylogeny-Guided Bayesian Framework for Reliable Microbiome Network Inference

**DOI:** 10.64898/2026.02.24.707538

**Authors:** Maryam Shahdoust, Rosa Aghdam, Golnaz Taheri

**Affiliations:** School of Biological Sciences, Institute for Research in Fundamental Sciences (IPM); Wisconsin Institute for Discovery, University of Wisconsin-Madison, Madison, WI, USA; School of Electrical Engineering and Computer Science, SciLifeLab, KTH Royal Institute of Technology, Stockholm, Sweden; School of Electrical Engineering and Computer Science, Digital Futures, KTH Royal Institute of Technology, Stockholm, Sweden

**Keywords:** Microbiome network inference, Bayesian Gaussian Graphical Model, Phylogenetic information, Precision matrix, Robustness

## Abstract

Understanding microbial interactions is essential for revealing the ecological structure and functional organization of microbiomes. However, network inference remains challenging due to the compositional, sparse, and high-dimensional nature of microbiome data, as well as the lack of gold-standard interaction benchmarks. Here, we introduce `PhyMapNet`, a Bayesian Gaussian Graphical Model framework that explicitly integrates phylogenetic information to infer microbiome networks. We introduce `PhyMapNet`, a phylogeny-aware method for reconstructing microbial association networks. `PhyMapNet` infers conditional dependencies among taxa by estimating a sparse precision matrix, while incorporating evolutionary relatedness through a kernel defined on phylogenetic distances. This integration of biological prior information improves the robustness and interpretability of the inferred networks. We evaluated `PhyMapNet` on two real-world microbiome datasets (Smoking and Caffeine), performing extensive robustness analyses under bootstrap and noisy perturbations, which demonstrated the reproducibility of the algorithm. The results also revealed that network structure can vary with parameter selection, highlighting the algorithm’s sensitivity to hyperparameters. Because `PhyMapNet` is computationally efficient and can explore thousands of parameter configurations within an hour, we developed a tuning-free framework that constructs reliable consensus networks by aggregating results across a broad hyperparameter space. This strategy yields consensus microbiome networks with controllable sparsity, offering a practical way to adjust network density while retaining highly stable edges. We then compared these consensus networks with those inferred by nine established methods and observed statistically significant overlap. To support reproducibility and practical use, we provide an open-source `R` package implementation of `PhyMapNet`.

## Introduction

Microbial communities are central to human health, agriculture, and ecosystem function, yet their complexity makes it difficult to uncover the interactions that shape their dynamics [1]. A widely used strategy to study these communities is the construction of microbiome networks, where nodes represent microbial taxa and edges represent inferred associations or interactions [2, 3]. Such networks provide valuable insights into ecological dependencies, identify candidate keystone species, and highlight potential biomarkers of disease or environmental change [4, 5, 6].

Constructing reliable microbial networks poses a major challenge, as the sparsity, compositional nature, and high dimensionality of microbiome data can obscure true biological relationships and yield inconsistent results across studies [7, 5]. Numerous algorithms have been developed to mitigate these issues, including correlation-based methods such as `SparCC` [8] and `CClasso` [9], as well as conditional-dependence approaches such as `SPIEC-EASI` [10]. However, the networks inferred by these tools often show limited overlap, reflecting differences in underlying statistical assumptions and raising concerns about the robustness of identified associations [11, 6]. Unlike gene regulatory networks, which can occasionally be benchmarked against experimentally validated pathways [12, 13], no universally accepted gold standard exists for microbiome interactions. This absence of a reference makes it difficult to evaluate method accuracy, leading to inconsistent ecological interpretations across studies [5, 14]. These limitations highlight the need for biologically informed network inference approaches that leverage evolutionary relationships to enhance robustness and interpretability.

Incorporating phylogenetic information has become increasingly important in microbiome research, as evolutionary distances provide biological context that abundance data alone cannot capture [15]. Foundational metrics such as `UniFrac` established phylogeny-based beta-diversity as a gold standard for distinguishing microbial community differences across environments [16]. Building on this principle, the `PhILR` transform integrates tree structure into compositional data analysis, reducing statistical artifacts and yielding more stable representations of microbial dynamics [17]. More recent network-focused frameworks explicitly embed evolutionary distances into inference. For example, `PGLasso` incorporates phylogenetic distances as priors in a graphical lasso model to improve the accuracy of microbial association networks [18], while `PhyloMint` adjusts metabolic competition and complementarity indices by evolutionary distance to construct ecologically meaningful microbial networks [19]. Beyond these methods, recent reviews emphasize that leveraging phylogenetic trees enhances statistical power, accounts for the non-independence of related taxa, and improves interpretability in microbiome analysis [20, 21]. Collectively, these advances demonstrate that phylogenetic tree information is both biologically meaningful and statistically advantageous across microbiome studies, motivating its use in network inference to generate more robust and ecologically grounded interaction maps.

Gaussian graphical models (GGMs) are widely applied in modern biological research to infer networks from high-dimensional data, as they estimate conditional dependencies (partial correlations) while accounting for uncertainty and sparsity [22]. This property makes GGMs particularly useful for inferring sparse networks from high-dimensional data [23, 24]. However, direct application of classical GGMs to microbiome sequencing data is problematic, since these data are zero-inflated, compositional, and heavily skewed, which violates Gaussian assumptions and leads to spurious associations. To address these challenges, several extensions have been developed, including latent Gaussian copula models, zero-inflated graphical models, and compositionality-aware approaches such as `SPIEC-EASI` [25] and `SPRING` [26], which adapt the GGM framework to microbiome-specific characteristics. These advances demonstrate the flexibility of GGMs for microbiome research while highlighting the need for further biologically informed extensions.

In this work, we introduce a novel Bayesian GGM framework, Phylogenetic Tree–Guided Mapping Network (`PhyMapNet`), that explicitly integrates phylogenetic information into microbiome network inference. Our approach utilizes kernel functions derived from phylogenetic distances to guide network estimation, enabling the incorporation of evolutionary relationships in a principled and scalable manner. `PhyMapNet` integrates statistical dependencies with evolutionary context, using phylogenetic information as external knowledge about relationships among taxa in microbiome studies. A key characteristic of `PhyMapNet` is its parameter-free design, which accounts for edge reliability and facilitates the inference of robust microbiome networks. To evaluate the performance of the proposed framework, we applied `PhyMapNet` to two distinct microbiome datasets: the Smoking dataset [27] and the Caffeine dataset [28]. These datasets span different microbial environments and provide a diverse test bed for validating the robustness and biological relevance of the constructed networks. By leveraging phylogenetic relationships and a Bayesian foundation, `PhyMapNet` delivers more stable and interpretable microbiome networks. We provide an open-source R package that allows researchers to apply `PhyMapNet` to microbiome data and corresponding phylogenetic trees, enabling efficient and reproducible inference of microbial interaction networks https://github.com/rosaaghdam/PhyMapNet.

## Methods

### PhyMapNet Algorithm

`PhyMapNet` employs Bayesian GGMs [29, 30] to infer conditional dependence between microbial taxa. The framework combines statistical dependencies with evolutionary distances derived from phylogenetic trees, yielding biologically grounded microbiome networks. The input of `PhyMapNet` consists of a microbiome abundance table of dimension *N × P*, where *N* is the number of samples and *P* is the number of taxa in the raw dataset. After filtering low-quality samples and rare taxa, the resulting abundance matrix has dimension *n × p* where *n* ≤ *N* and *p* ≤ *P*. In addition to the filtered abundance matrix, the corresponding phylogenetic tree containing the same *p* taxa serves as an input to the algorithm. `PhyMapNet` proceeds through a series of steps to construct microbiome networks from these inputs. Figure 1 provides a graphical abstract of the algorithmic workflow for Steps 1–4, illustrating the process from preprocessing to network construction based on a single set of parameters. In Step 5, the final reliable network is constructed by integrating multiple networks obtained from different parameter settings.

**Fig. 1.**
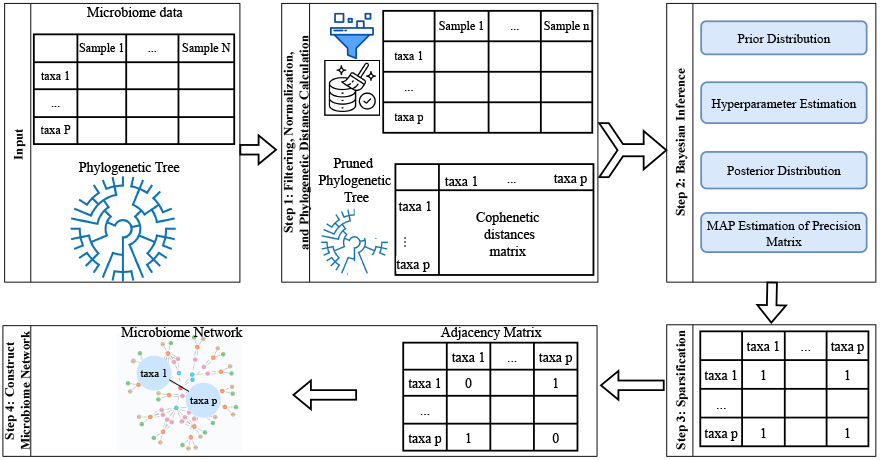
Overview of the PhyMapNet workflow. **Step 1:** Microbiome count data are filtered and normalized, and phylogenetic distances between taxa are computed from a pruned phylogenetic tree.**Step 2:** A Bayesian inference framework incorporates phylogenetic information through structured prior distributions on model parameters.**Step 3:** Sparsification is applied to the estimated precision matrix using a maximum a posteriori (MAP) criterion to encourage network sparsity. **Step 4:** The microbiome network is constructed by converting the sparse precision matrix into a binary adjacency matrix representing conditional dependencies among taxa. In the reliability framework, this entire procedure is repeated across multiple hyperparameter configurations to assess edge stability and construct consensus networks.

### Step 1: Filtering, Normalization, and Phylogenetic Distance Calculation

#### Filtering

Samples containing fewer than 15 observed (nonzero) taxa were excluded. Taxa with zero counts in more than a predefined proportion of samples (90% for the Smoking dataset and 75% for the Caffeine dataset) were removed. Additionally, taxa with a median nonzero abundance of zero across samples were excluded, ensuring that only taxa with meaningful prevalence and abundance were retained for downstream analysis. After filtering, the *n × p* microbiome abundance table was constructed.

#### Normalization

Raw count data were normalized to mitigate compositional effects, sequencing depth variations, and other technical artifacts that can bias network inference. Normalization has been shown to strongly affect the sparsity and accuracy of inferred networks [31]. We evaluated four widely used normalization strategies implemented in our framework: (1) Log transformation, log(count + 1), which stabilizes variance and reduces the influence of highly abundant taxa [32]; (2) GMPR (Geometric Mean of Pairwise Ratios), which corrects for library size differences and compositional bias [33]; (3) CLR (Centered Log-Ratio) transformation, which accounts for the compositional structure of microbiome data by scaling counts relative to the geometric mean across taxa after adding a small pseudo-count (0.01) [25]; and (4) TSS (Total Sum Scaling), which normalizes each sample by its total read count and optionally rescales to a common factor.

#### Phylogenetic distances

To integrate evolutionary context, pairwise cophenetic distances between taxa were computed from the pruned phylogenetic tree [34]. This step used the `ape::cophenetic.phylo` function in `R` [35] to produce a symmetric *p × p* distance matrix fully aligned with the *n × p* filtered and normalized taxa table, ensuring that the evolutionary structure corresponds exactly to the retained taxa.

### Step 2: Bayesian Inference

`PhyMapNet` employs GGMs [29] within a Bayesian inference framework to learn conditional dependencies among microbial taxa from their joint distribution, assuming multivariate normality of the normalized abundance data. Specifically, let *Y*_*s*_ ∼ 𝒩_*p*_(0, Θ^−1^) for *s* = 1, …, *n*, where *Y*_*s*_ ∈ ℝ^*p*^ denotes the normalized abundance vector for sample *s* and Θ ∈ ℝ^*p×p*^ denotes the precision matrix encoding conditional dependencies among taxa. In this framework, the precision matrix Θ encodes the relationships among microbial taxa: non-zero off-diagonal elements indicate conditional dependencies. The partial correlation between taxa *i* and *j* is given by 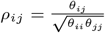, where *θ*_*ij*_ is the element of precision matrix Θ. The likelihood function of *Y*_*s*_ ∼ *N* (0, Θ^−1^) is:

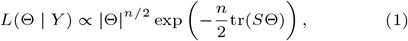

where *S* is the sample covariance matrix. To estimate the precision matrix Θ, `PhyMapNet` adopts a Bayesian framework by placing a Wishart prior *W* (*ν, G*) on Θ. The prior parameter *G* = (*ν*Ω)^−1^, where Ω incorporates phylogenetic information. Phylogenetic distances are introduced through kernel functions [36, 37, 38], which transform cophenetic distances between taxa into a covariance structure reflecting prior biological knowledge. The prior hyperparameter is specified as

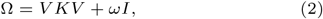

where *V* = Diag(*σ*_1_, *σ*_2_, …, *σ*_*p*_) is a *p × p* diagonal matrix of microbial standard deviations, *K* is a kernel matrix derived from phylogenetic distances, and *ωI* ensures positive definiteness. In this study, we considered two stationary kernels [39]:

1. Squared Exponential (SE) kernel [40, 41, 42], also known as Gaussian or Radial Basis Function (RBF) kernel:

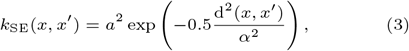
2. Ornstein–Uhlenbeck (OU) kernel [42], a member of the Matérn family:

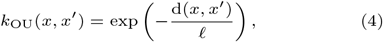

where d(*x, x*^*′*^) is the cophenetic distance between taxa. In equation 3, *a*^2^ is the signal variance (often set to 1), and *α* is the length scale controlling smoothness. In equation 4, ℓ controls the rate of correlation decay with the phylogenetic distance. The degrees of freedom parameter *ν* is another hyperparameter of the Wishart prior. It can be set empirically as any non-negative real number greater than *p* − 1, with initial value *p* + 1 [43, 44]. Following recommendations by Zhang et al. [45], we set *ν* = 2*p*, where *p* is the number of taxa retained after filtering. This choice reflects the strength of belief in the prior hyperparameters. Some literature suggests setting *ν >* 2*p* [45, 43, 44].

The posterior distribution of Θ is also Wishart with updated parameters:

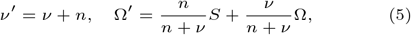

as shown in [46, 47].

Finally, the maximum a posteriori (MAP) estimator of the precision matrix is

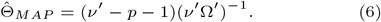

Two additional hyperparameters, *ϵ*_1_ and *ϵ*_2_, are introduced as regularization terms to ensure numerical stability and positive definiteness during Bayesian estimation. Specifically, *ϵ*_1_ is added to the diagonal of the kernel-based prior covariance Ω to stabilize its inversion, while *ϵ*_2_ is applied to the posterior precision estimate to prevent singularity in the resulting precision matrix. Both parameters act as small positive constants that mitigate computational instability without materially affecting the inferred conditional dependencies.

### Step 3: Sparsification

The posterior estimation of the precision matrix is the mode of the MAP, which does not have zero components. However, for microbiome networks, sparsity is essential to highlight meaningful conditional dependencies and reduce spurious associations. Therefore, to impose sparsity on the MAP, the hard-thresholding method is applied [48]. In this process, we applied percentile-based cutoffs on the absolute values of the estimated conditional correlations in the MAP. Edges with values below the chosen threshold (*th*_sparsity_) were set to zero, yielding a sparse precision matrix that defines the final microbiome network.

### Step 4: Infer microbiome network

The adjacency matrix (*AD*) is constructed from the estimated sparse matrix in the following manner:

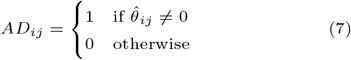

where *i, j* ∈ {1, …, *p*} and *p* is the number of microbial taxa. The non-zero elements indicate conditional dependencies, which are represented as edges in the final microbiome network.

### Step 5: Construct the final reliable microbiome network

The reliability of inferred microbial associations was evaluated through an extensive hyperparameter ensemble approach designed to assess the robustness of network edges across a wide range of parameter settings. This procedure produces an empirical measure of edge stability, mitigating sensitivity to arbitrary hyperparameter choices, and reducing the risk of overfitting to any specific configuration. To comprehensively assess edge stability, we explored a broad hyperparameter space ℋ. The ensemble grid systematically varied four key parameters: kernel bandwidth *α*, neighborhood size *k*, and regularization terms *ϵ*_1_ and *ϵ*_2_. Each combination was evaluated under a fixed sparsification threshold (*th*_sparsity_), enforcing strong network sparsity. For the kernel function, two options were considered, OU and Squared SE kernels, both of which integrate phylogenetic distances into the covariance structure. For each configuration, the data were normalized using one of four common normalization strategies, log transformation, GMPR, CLR, or TSS. Together, this parameter grid defines the complete hyperparameter ensemble ℋ, ensuring that the resulting reliability scores reflect robustness across realistic and biologically plausible model variations.

#### Reliability score

Each run *h* represents a distinct hyperparameter configuration from the ensemble ℋ and produces a binary adjacency matrix *AD*^(*h*)^ ∈ {0, 1}^*p×p*^. For every potential edge (*i, j*), the reliability score was computed as:

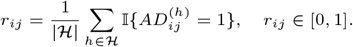

Here, ℋ denotes the complete set of hyperparameter configurations considered in the ensemble, and |*H*| is the total number of runs (approximately 10^4^ in this study). A value of *r*_*ij*_ = 1 indicates that edge (*i, j*) was consistently recovered across all runs whereas *r*_*ij*_ = 0 means it never appeared. This ensemble-based measure represents the relative frequency of detection of each edge, providing a robust, data-driven estimate of edge stability. The resulting outputs comprise (i) a weighted reliability matrix *R* = (*r*_*ij*_), where each element *r*_*ij*_ represents the relative frequency of detection of edge (*i, j*), and (ii) a binary consensus network obtained by thresholding *R* at *r*_*ij*_ ≥ *th*_reliability_, typically *th*_reliability_ = 0.7, retaining only edges with high reliability across models.

The symbols and parameters used throughout the `PhyMapNet` framework and evaluation protocol are summarized in Table 1.

**Table 1.**
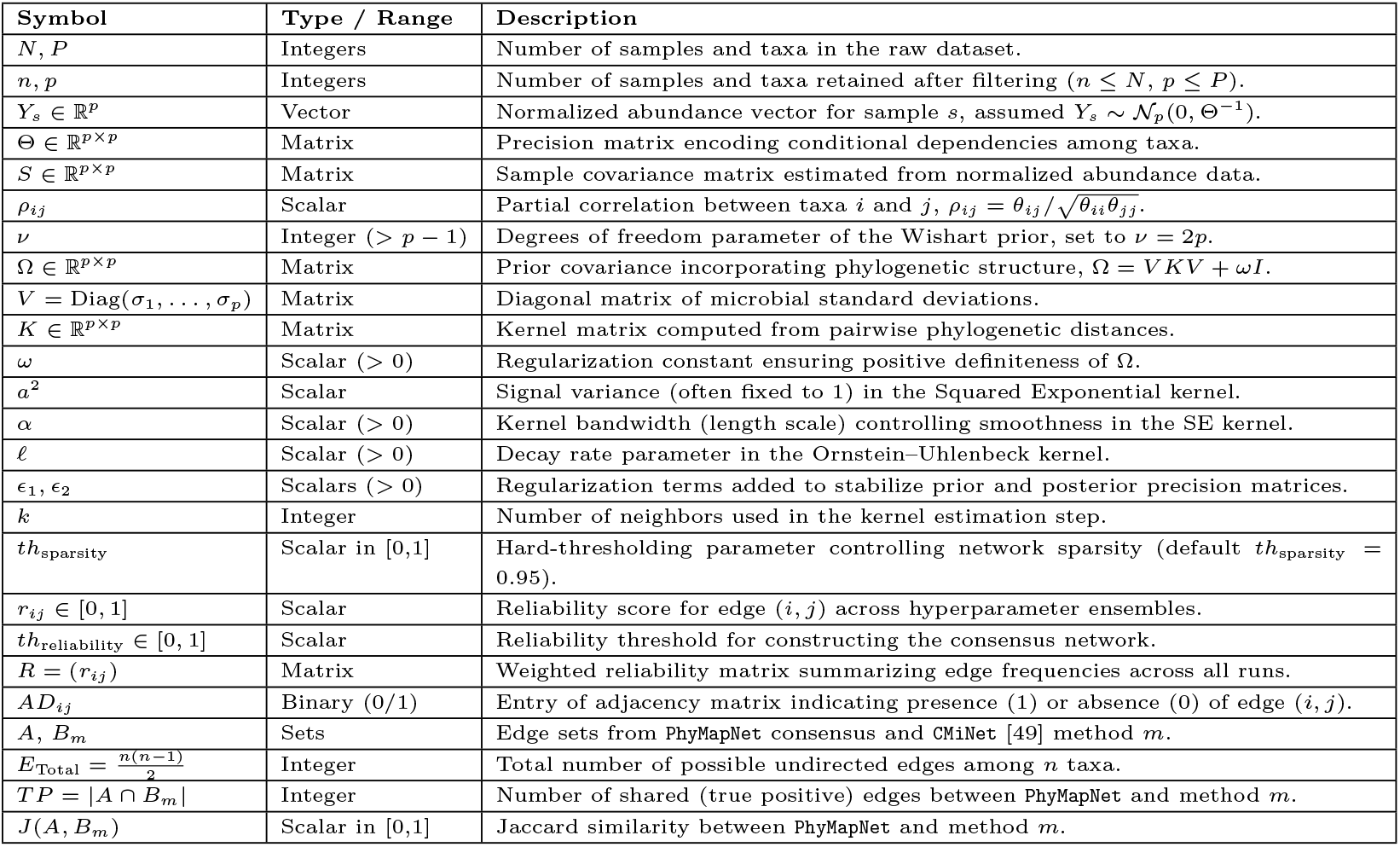
Summary of notation and parameters used in the PhyMapNet framework and evaluation protocol.

### Evaluation protocol

We evaluate `PhyMapNet` using (i) robustness analyses under bootstrap resampling and noise perturbations, (ii) a reliability-based consensus construction across hyperparameter settings, and (iii) concordance with networks inferred by established methods.

#### Robustness analysis

Because no gold-standard microbial network is available for real-world datasets, direct evaluation of biological accuracy is not feasible. Therefore, we evaluated the robustness of `PhyMapNet` to data perturbations as an indirect measure of methodological stability and reproducibility. This analysis evaluates whether the inferred network structure remains consistent under realistic variations in the input data, rather than whether it reflects the true underlying biological relationships. To assess stability, we performed two complementary robustness evaluations for each dataset. (1) *Noisy datasets*. We generated 100 noisy datasets by perturbing a random subset of abundance values in each sample using a truncated normal distribution with mean and variance matched to the observed data. The added noise was constrained within the empirical range of observed values to preserve biological realism. (2) *Bootstrap datasets*. We generated 100 bootstrap replicates by resampling samples with replacement to capture variability arising from sample composition. For both analyses, each perturbed dataset was processed independently through the full `PhyMapNet` workflow, and the resulting networks were compared to the reference network (constructed from the original data) using the F1-score. In addition to evaluating `PhyMapNet`, we included `SPIEC-EASI` [25] as a benchmark method for comparison, as it similarly infers microbial association networks through precision matrix estimation implemented through the graphical lasso [50]. To ensure fairness, the sparsity level of `PhyMapNet` was calibrated to yield a comparable number of edges to those produced by `SPIEC-EASI` under its default settings, thereby isolating structural differences attributable to inference methodology rather than network density.

To quantify uncertainty in network reconstruction, 95% bootstrap confidence intervals [51] were computed for the F1-scores, which measure how similar the reconstructed networks are to the reference network obtained from the original data. A narrow confidence interval indicates that small changes in the data produce nearly identical network structures, demonstrating the reproducibility of the algorithm. In contrast, a wide interval would suggest that the inferred networks vary substantially across resampled datasets. Moreover, higher lower bounds of the confidence interval correspond to consistently high F1-scores, indicating that even under data perturbations, the reconstructed networks remain highly similar to the network inferred from the original data, reflecting the reliability of the algorithm under noise and resampling.

#### Reliability-based consensus network

We quantified edge stability using a large hyperparameter ensemble (Step 5 in the `PhyMapNet` Algorithm). Across ≈ runs per dataset, we systematically varied the kernel bandwidth

(*α*), neighborhood size (*k*), and model parameters (*ϵ*_1_, *ϵ*_2_) over a predefined grid, combined with four data normalizations (log, GMPR, CLR, TSS) and kernels (SE or OU). Each model produced an adjacency matrix after applying a fixed sparsification threshold to the precision matrix (*th*_sparsity_), reflecting the biological prior of strong network sparsity. For each potential edge (*i, j*), the reliability score *r*_*ij*_ is the empirical frequency of detection across runs in the ensemble in large hyperparameter ensemble. The resulting reliability matrix *R* = (*r*_*ij*_) summarizes edge stability across the full parameter range.

We defined a final *consensus* microbiome network by thresholding *R* at a reliability cutoff *th*_reliability_, retaining only edges with *r*_*ij*_ ≥ *th*_reliability_. Unless stated otherwise, *th*_reliability_ was chosen to achieve a sparsity level comparable to that of other algorithms in the `CMiNet` package (e.g., *th*_reliability_ = 0.65 for Smoking and *th*_reliability_ = 0.55 for Caffeine).

#### Concordance with existing methods

To assess concordance with widely used approaches, we compared the `PhyMapNet` consensus network with the nine algorithms implemented in the `CMiNet` package (Pearson [3], Spearman [52], biweight midcorrelation [4], `SparCC` [8], `SPIEC-EASI` (MB and glasso variants) [25], `SPRING` [26], `CClasso` [9], and `CMIMN` [53]).

##### Statistical significance of overlap

Let *A* denote the set of edges in the `PhyMapNet` consensus and *B*_*m*_ the edges of the CMiNet method *m*. We report: (i) the number of edges in *A* (|*A*|) and in each *B*_*m*_ (|*B*_*m*_|); (ii) the size of the intersection |*A* ∩ *B*_*m*_|; (iii) the Jaccard similarity:

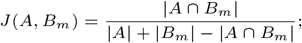

and (iv) we computed a one-tailed hypergeometric *p*-value to assess whether the observed overlap between the `PhyMapNet` network and each individual method could occur by chance:

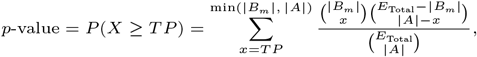

where *TP* = |*A* ∩ *B*_*m*_| and 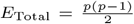 is the number of possible undirected edges among *p* taxa. In practice, *P* (*X* ≥ *TP*) was computed using the `phyper` function in R with `lower.tail = FALSE`.

## Results

### Datasets

The Smoking dataset [27] analyzed in this study originates from research examining the effects of smoking on the human upper respiratory tract microbiome. The dataset comprises an Operational Taxonomic Unit (OTU) abundance matrix, metadata on smoking status, and a phylogenetic tree. The dataset underwent preprocessing to remove low-coverage samples and sparse taxa while ensuring consistency with the phylogenetic tree. Specifically, samples with fewer than 15 observed (nonzero) taxa were removed to ensure sufficient microbial representation. OTUs with zero counts in more than 90% of samples were excluded based on a zero-proportion threshold of 0.90, and OTUs whose nonzero median abundance across samples was zero were also discarded. After filtering, the retained OTUs were aligned with the phylogenetic tree tips to ensure one-to-one correspondence, and the tree was pruned accordingly. The final dataset contained 60 samples and 174 OTUs with nonzero median abundance. Phylogenetic distances between OTUs were computed from the pruned tree using the `ape::cophenetic.phylo` function [35], producing a symmetric *p × p* cophenetic distance matrix aligned with the filtered OTU table.

The Caffeine dataset was downloaded from Qiita (https://qiita.ucsd.edu/, study ID 1,011) and contains 98 samples and 6,674 OTUs. Caffeine intake was selected as the outcome of interest, as caffeine consumption has been shown to significantly influence gut microbiota composition. The dataset includes an OTU abundance matrix, metadata [28] caffeine intake levels, and a corresponding phylogenetic tree. A series of preprocessing steps was applied to ensure data quality and consistency with the phylogenetic tree. Samples with fewer than 15 observed taxa were removed, and OTUs with zero counts in more than 75% of samples (zero-proportion threshold = 0.75) were excluded. OTUs with a median nonzero abundance of zero were also discarded. After filtering, the final dataset contained 98 samples and 299 OTUs. The phylogenetic tree was pruned to retain only tips corresponding to the filtered OTUs, and the cophenetic (patristic) distance matrix was computed from the pruned tree using `ape::cophenetic.phylo` to quantify pairwise evolutionary distances among OTUs. Table 2 summarizes the number of samples and OTUs before and after filtering, the zero-count threshold applied to each dataset, and their corresponding sources.

**Table 2.**
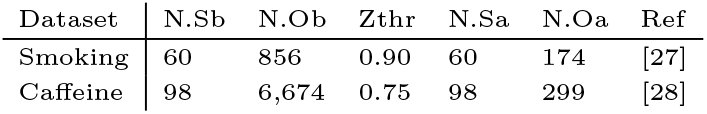
Summary of datasets and preprocessing. N.Sb = number of samples before filtering; N.Ob = number of OTUs before filtering; Zthr = zero-count threshold; N.Sa = number of samples after filtering; N.Oa = number of OTUs after filtering.

### Results of robustness analysis

To comprehensively evaluate robustness, we explored a range of hyperparameter configurations encompassing different values of *α*, kernel type, *k, ϵ*_1_, and *ϵ*_2_. The resulting F1-scores spanned a broad range (approximately 0.6–1.0), reflecting the sensitivity of network inference to model parameterization. Rather than exhaustively searching all possible combinations, we selected two representative configurations, `PhyMapNet1` and `PhyMapNet2`, to illustrate the variability in performance across this range. `PhyMapNet1` corresponds to a high-performing configuration (*α* = 0.1, Laplacian kernel, *k* = 10, *ϵ*_1_ = 1, *ϵ*_2_ = 0.4), whereas `PhyMapNet2` represents a lower-performing but stable configuration (*α* = 0.01, Laplacian kernel, *k* = 5, *ϵ*_1_ = *ϵ*_2_ = 0.5).

For each dataset, the sparsity level of `PhyMapNet` was calibrated to yield a comparable number of edges to those obtained by `SPIEC-EASI` under its default parameters, ensuring similar network densities for a fair baseline comparison of stability. In the networks inferred from the original data, `SPIEC-EASI` produced 269 edges for the Smoking dataset and 530 edges for the Caffeine dataset. After tuning, the final `PhyMapNet` configurations generated 302 edges (Smoking) and 446 edges (Caffeine), providing closely matched levels of sparsity. All other hyperparameters were held constant across datasets, except for the sparsification threshold (*th*_sparsity_), which was adjusted to align the overall edge density (*th*_sparsity_ = 0.98 for Smoking and *th*_sparsity_ = 0.99 for Caffeine). These edge counts correspond to the reference networks inferred from the original data, which were used for evaluating robustness and calculating F1-scores across bootstrap and noisy perturbations.

Table 3 summarizes the F1-score distributions across 100 bootstrap and noisy replicates for both datasets and all three methods, each evaluated against its respective reference network. The table reports comprehensive summary statistics, including the mean, standard deviation, minimum, maximum, quartiles (Q25, Q50, Q75), and 95% confidence interval bounds (CI l and CI u), along with CI r, representing the interval width. Narrower CI r values and smaller standard deviations indicate higher reproducibility and stability, while wider intervals reflect increased sensitivity to data perturbations. Overall, `PhyMapNet` shows consistently narrower confidence intervals and smaller variability than `SPIEC-EASI`, confirming its robustness and stability under both bootstrap and noisy resampling conditions.

**Table 3.** Robustness of `PhyMapNet1`, `PhyMapNet2`, and `SPIEC-EASI` under bootstrap and noisy resampling for the Smoking and Caffeine datasets. `PhyMapNet1` (*α* = 0.1, Laplacian kernel, *k* = 10, *ϵ*_1_ = 1, *ϵ*_2_ = 0.4) and `PhyMapNet2` (*α* = 0.01, Laplacian kernel, *k* = 5, *ϵ*_1_ = *ϵ*_2_ = 0.5) represent two illustrative parameter configurations corresponding to different ranges of F1-score values. Each entry summarizes the distribution of F1-scores obtained from 100 perturbed datasets and compared with the reference network inferred from the original data. Reported statistics include the mean (ME), standard deviation (SD), minimum (Min), maximum (Max), quartiles (Q25, Q50, Q75), and the lower (CI l) and upper (CI u) bounds of the 95% confidence interval, as well as the CI r, representing the interval width.

|  | Method | Split | ME | SD | Min | Max | Q25 | Q50 | Q75 | CI<sub>l</sub> | CI<sub>u</sub> | CI<sub>r</sub> |
| --- | --- | --- | --- | --- | --- | --- | --- | --- | --- | --- | --- | --- |
| Smoking | PhyMapNet1 | Bootstrap | 0.94 | 0.01 | 0.92 | 0.97 | 0.94 | 0.94 | 0.95 | 0.92 | 0.96 | 0.04 |
|  | PhyMapNet1 | Noisy | 0.96 | 0.01 | 0.94 | 0.98 | 0.96 | 0.96 | 0.97 | 0.95 | 0.97 | 0.02 |
|  | PhyMapNet2 | Bootstrap | 0.67 | 0.03 | 0.60 | 0.74 | 0.65 | 0.67 | 0.69 | 0.62 | 0.73 | 0.10 |
|  | PhyMapNet2 | Noisy | 0.69 | 0.02 | 0.65 | 0.73 | 0.68 | 0.69 | 0.70 | 0.66 | 0.72 | 0.06 |
|  | SPIEC-EASI | Bootstrap | 0.52 | 0.05 | 0.37 | 0.65 | 0.49 | 0.52 | 0.55 | 0.42 | 0.62 | 0.20 |
|  | SPIEC-EASI | Noisy | 0.55 | 0.03 | 0.46 | 0.62 | 0.53 | 0.55 | 0.57 | 0.49 | 0.60 | 0.11 |
| Caffeine | PhyMapNet1 | Bootstrap | 0.97 | 0.01 | 0.96 | 0.98 | 0.97 | 0.97 | 0.97 | 0.96 | 0.98 | 0.02 |
|  | PhyMapNet1 | Noisy | 0.98 | 0.00 | 0.97 | 0.99 | 0.98 | 0.98 | 0.98 | 0.97 | 0.99 | 0.01 |
|  | PhyMapNet2 | Bootstrap | 0.61 | 0.05 | 0.51 | 0.73 | 0.59 | 0.62 | 0.64 | 0.53 | 0.70 | 0.17 |
|  | PhyMapNet2 | Noisy | 0.67 | 0.02 | 0.61 | 0.72 | 0.66 | 0.67 | 0.69 | 0.64 | 0.71 | 0.07 |
|  | SPIEC-EASI | Bootstrap | 0.49 | 0.04 | 0.38 | 0.57 | 0.47 | 0.50 | 0.52 | 0.42 | 0.57 | 0.15 |
|  | SE<sub>glasso</sub> | Noisy | 0.58 | 0.02 | 0.50 | 0.63 | 0.57 | 0.59 | 0.60 | 0.51 | 0.61 | 0.10 |

Figure 2 visualizes the distribution of F1-scores across these replicates for `PhyMapNet1`, `PhyMapNet2`, and `SPIEC-EASI`. Each box displays the variation in F1-score, with the 95% confidence interval shown above the box. The results show that both `PhyMapNet` configurations yield reproducible networks with F1-scores substantially higher than those of `SPIEC-EASI`, underscoring the algorithm’s robustness across parameter variations and data perturbations. However, because the true microbial associations are unknown, these results primarily demonstrate reproducibility rather than biological accuracy.

**Fig. 2.**
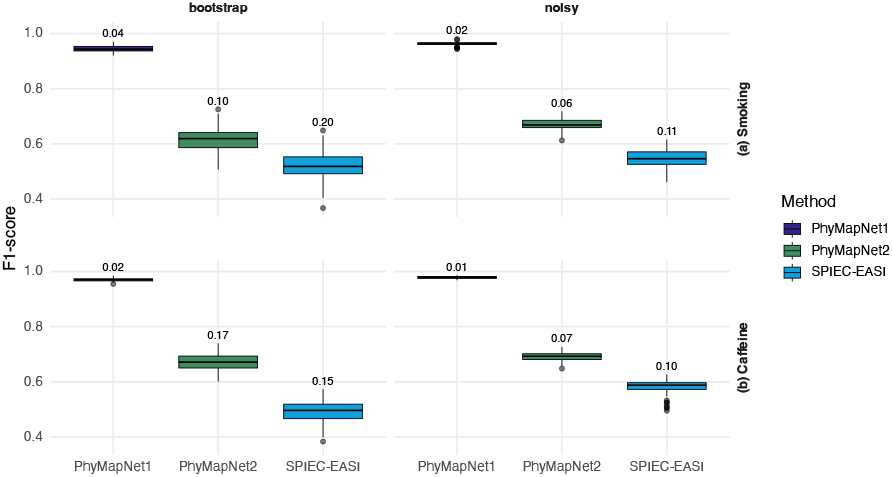
Distribution of F1-scores under bootstrap and noisy resampling for `PhyMapNet1`, `PhyMapNet2`, and `SPIEC-EASI` across the (a) Smoking and (b) Caffeine datasets. `PhyMapNet1` (*α* = 0.1, Laplacian kernel, *k* = 10, *ϵ*_1_ = 1, *ϵ*_2_ = 0.4) and `PhyMapNet2` (*α* = 0.01, Laplacian kernel, *k* = 5, *ϵ*_1_ = *ϵ*_2_ = 0.5) represent two illustrative parameter configurations corresponding to different ranges of F1-score values. Each boxplot represents the distribution of F1-scores obtained from 100 replicate networks reconstructed from perturbed datasets. Numerical values shown above each box correspond to the 95% confidence interval (CI) width for the respective F1-score distribution.

### Reliability-driven consensus and external support

To comprehensively assess edge stability, we explored a broad hyperparameter space *H* consisting of approximately 10^4^ model configurations. The ensemble grid systematically varied four key parameters: kernel bandwidth *α* ∈ [0.01, 0.12] (step size 0.01), neighborhood size *k* ∈ [2, 10], and regularization terms *ϵ*_1_, *ϵ*_2_ ∈ [0, 1] (step size 0.1). Figure 3 illustrates the reliability of external support for each common edge (*i, j*) by plotting its reliability score *r*_*ij*_ (x-axis) against the aggregated `CMiNet` support score (y-axis; 0–9 methods). This value indicates how many of the nine algorithms confirmed the presence of that edge. For example, a score of 0 means the edge was not detected by any method, 1 means it was supported by one method, and 9 means it was consistently inferred by all nine methods. X-axis ticks show bin ranges with counts of edges per bin. Edges above the reliability threshold *th*_reliability_ are highlighted by blue color. Using reliability thresholds chosen to match network sparsity to other nine algorithms, the `PhyMapNet` consensus networks retained a subset of highly stable edges (Smoking: *th*_reliability_ = 0.65, yielding 216 edges; Caffeine: *th*_reliability_ = 0.55, yielding 687 edges). Across both datasets, edges with higher reliability were more frequently supported by multiple `CMiNet` methods (Figure 3), indicating a strong positive association between stability under hyperparameter variation and external methodological agreement. Interestingly, no edges with low reliability were supported by many algorithms (particularly when the number of supporting methods exceeds six), suggesting that unstable associations are less reproducible across methods. At the same time, several unique edges identified by `PhyMapNet` were not detected by any other method. Such distinctive associations are common in microbiome network studies and may reflect biologically meaningful relationships that remain undetected by other approaches.

**Fig. 3.**
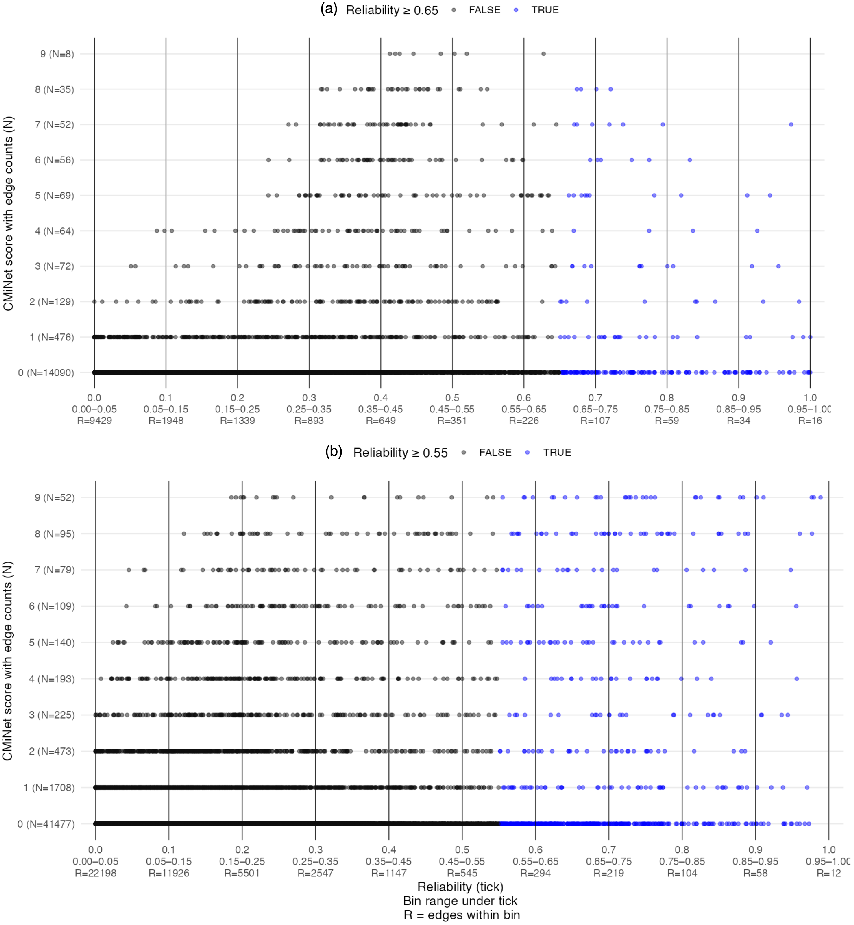
Reliability (*r*_*ij*_) versus aggregated `CMiNet` support (0–9 methods) for all edges on the shared taxon universe: (a) Smoking and (b) Caffeine. X-axis ticks show the bin centers; the second line under each tick indicates the bin range; “R=” gives the number of edges within the bin. Points above the dataset-specific reliability threshold *th*_reliability_ are highlighted. Higher reliability is associated with greater external support across methods.

For each of the nine `CMiNet` algorithms, we quantified the overlap with the `PhyMapNet` consensus network, reporting the number of overlapping edges, total edges per method, Jaccard similarity, and one-tailed hypergeometric *p*-values (Table 4). Multiple methods showed statistically significant enrichment of shared edges (p−*value* ≪ 0.05), and several achieved modest Jaccard similarities with `PhyMapNet`.

**Table 4.** Overlap statistics between the `PhyMapNet` consensus network and each of the nine `CMiNet` algorithms across the Smoking and Caffeine datasets. The `PhyMapNet` consensus networks contained 216 edges for Smoking and 687 edges for Caffeine. For each method, the number of overlapping edges, total number of inferred edges, percentage of `PhyMapNet` edges confirmed, Jaccard similarity, and one-tailed hypergeometric *p*-value are reported. Multiple methods exhibit statistically significant enrichment of shared edges (*p*-value ≪ 0.05), with generally modest Jaccard values (≈ 0.03–0.16), reflecting partial but non-random agreement between `PhyMapNet` and existing inference approaches.

|  | Method | Overlap | Edges (Method) | % of PhyMapNet | Jaccard | p-value |
| --- | --- | --- | --- | --- | --- | --- |
| Smoking | bicor | 34 | 216 | 15.74 | 0.085 | 8.48E–26 |
|  | SPIEC-EASI_mb | 32 | 205 | 14.81 | 0.082 | 3.39E–24 |
|  | SPIEC-EASI_glasso | 47 | 450 | 21.76 | 0.076 | 1.99E–27 |
|  | SPRING | 51 | 565 | 23.61 | 0.070 | 8.72E–27 |
|  | CMiMN | 29 | 303 | 13.43 | 0.059 | 4.68E–16 |
|  | pearson | 19 | 216 | 8.80 | 0.046 | 2.97E–10 |
|  | spearman | 19 | 216 | 8.80 | 0.046 | 2.97E–10 |
|  | SparCC | 16 | 216 | 7.41 | 0.038 | 8.99E–08 |
|  | CClasso | 14 | 216 | 6.48 | 0.033 | 2.89E–06 |
| Caffeine | spearman | 195 | 745 | 28.38 | 0.158 | 2.75E–185 |
|  | CMiMN | 212 | 895 | 30.86 | 0.155 | 2.05E–192 |
|  | CClasso | 174 | 745 | 25.33 | 0.138 | 9.03E–155 |
|  | spiecEasi_glasso | 108 | 316 | 15.72 | 0.121 | 1.39E–114 |
|  | spiecEasi_mb | 126 | 587 | 18.34 | 0.110 | 1.28E–105 |
|  | SparCC | 134 | 745 | 19.51 | 0.103 | 9.24E–102 |
|  | pearson | 131 | 745 | 19.07 | 0.101 | 4.47E–98 |
|  | bicor | 127 | 745 | 18.49 | 0.097 | 3.14E–93 |
|  | SPRING | 187 | 1713 | 27.22 | 0.085 | 3.72E–104 |

Because no gold-standard microbial network is available, these analyses assess *reproducibility and external concordance* rather than biological ground truth. Taken together, the observed patterns of reliability, `CMiNet` support, and overlap enrichment indicate that the reliability-filtered edges identified by `PhyMapNet` represent associations that are both stable under perturbations and broadly consistent with signals detected by diverse inference methods.

## Discussion

In this study, we introduced `PhyMapNet`, a Bayesian Gaussian Graphical Model framework that leverages phylogenetic information and precision-matrix estimation to infer microbiome networks. By integrating evolutionary distances through kernel functions, `PhyMapNet` embeds biological structure directly into network inference, providing a principled way to distinguish genuine ecological interactions from spurious correlations. Our extensive analyses across two real-world microbiome datasets (Smoking and Caffeine) demonstrated that `PhyMapNet` produces networks that are stable under bootstrap perturbations. In the absence of a gold-standard network, the narrow confidence intervals of the F1-scores indicate high reproducibility of the algorithm rather than direct biological accuracy. However, the bootstrap results based on two different parameter settings revealed that the inferred network structure can vary with parameter choice, reflecting the sensitivity of precision matrix–based inference to model hyperparameters. To address this limitation, `PhyMapNet` introduces a reliability-driven consensus strategy that aggregates information across thousands of parameter configurations to identify high-confidence, parameter-independent networks. This consensus framework enables users to focus on reproducible and consistent edges rather than results tied to a single parameter setting. Ultimately, by selecting a single global sparsity threshold, `PhyMapNet` constructs a reliable, parameter-free consensus network in which sparsity, and consequently the number of inferred edges, can be explicitly controlled.

The hyperparameter ensemble provides a parameter-free estimate of edge robustness for four key reasons: (1) a fixed sparsification threshold (*th*_sparsity_) enforces strong sparsity consistent with biological constraints on microbial networks; (2) ensemble averaging removes dependency on specific hyperparameter settings, ensuring robustness across diverse configurations; (3) `PhyMapNet`’s computational efficiency allows rapid inference across thousands of models within practical runtime; and (4) in the absence of a gold-standard reference network, edge reliability offers a data-driven alternative to receiver operating characteristic (ROC)-based parameter tuning.

This represents a notable advantage over many existing methods that lack direct control over network density. A key factor enabling this framework is `PhyMapNet`’s computational efficiency, which allows more than 10,000 model evaluations to be completed in under one hour, a task that is infeasible for most Gaussian graphical model–based approaches. `PhyMapNet` achieves substantial computational efficiency because each model evaluation scales quadratically with the number of taxa, 𝒪 (*p*^2^). Phylogenetic distances are computed once and reused, and Bayesian posterior updates rely on closed-form matrix operations followed by element-wise sparsification. In contrast, Gaussian graphical model-based approaches typically require iterative optimization with cubic complexity 𝒪 (*p*^3^) per iteration due to repeated matrix inversions. As a result, `PhyMapNet` can perform more than 10,000 model evaluations within one hour, a scale that is computationally infeasible for most GGM-based methods. All runs completed comfortably on a workstation using 8 CPU cores and 16 GB of RAM. Benchmark comparisons against nine existing algorithms revealed significant overlap between `PhyMapNet` and other methods, while low-reliability edges were rarely confirmed by multiple approaches (fewer than six out of nine methods), highlighting the robustness of reliability-filtered associations. Nonetheless, the absence of experimentally validated ground-truth interactions remains a fundamental limitation, underscoring the need for integrative benchmarking and experimental validation in future work. `PhyMapNet` provides an open-source, scalable foundation for such efforts, facilitating the development of more reliable and biologically grounded frameworks for microbiome network analysis.

## Author contributions

M.SH. and R.A. conceived and designed the study. R.A. and M.SH. developed the `PhyMapNet` framework, implemented the software, and performed all computational experiments and data analysis. M.SH., R.A., and G.T. interpreted the results and contributed to methodological development. R.A. and M.SH. drafted the manuscript. G.T. supervised the research. All authors reviewed, edited, and approved the final manuscript.

## Conflict of interest

None declared.

## Funding

This work was supported by the SciLifeLab & Wallenberg Data Driven Life Science Program (grant: KAW 2024.0159).

## Data availability

The `PhyMapNet` source code is freely available at https://github.com/rosaaghdam/PhyMapNet.

## Notes

### Competing Interest Statement

The authors have declared no competing interest.

## References

1. Karoline Faust and Jeroen Raes. Microbial interactions: from networks to models. Nature Reviews Microbiology, 10(8):538–550, 2012.

2. Bing Guo, Lei Zhang, Huijuan Sun, Mengjiao Gao, Najiaowa Yu, Qianyi Zhang, Anqi Mou, and Yang Liu. Microbial co-occurrence network topological properties link with reactor parameters and reveal importance of low-abundance genera. npj Biofilms and Microbiomes, 8(1):3, 2022.

3. Monica Steffi Matchado, Michael Lauber, Sandra Reitmeier, Tim Kacprowski, Jan Baumbach, Dirk Haller, and Markus List. Network analysis methods for studying microbial communities: A mini review. Computational and structural biotechnology journal, 19:2687–2698, 2021.

4. Stefanie Peschel, Christian L Müller, Erika Von Mutius, Anne-Laure Boulesteix, and Martin Depner. Netcomi: network construction and comparison for microbiome data in r. Briefings in bioinformatics, 22(4):bbaa290, 2021.

5. Karoline Faust. Open challenges for microbial network construction and analysis. The ISME Journal, 15(11):3111–3118, 2021.

6. Kacie T Kajihara and Nicole A Hynson. Networks as tools for defining emergent properties of microbiomes and their stability. Microbiome, 12(1):184, 2024.

7. Zhaoqian Liu, Anjun Ma, Ewy Mathé, Marlena Merling, Qin Ma, and Bingqiang Liu. Network analyses in microbiome based on high-throughput multi-omics data. Briefings in bioinformatics, 22(2):1639–1655, 2021.

8. Jonathan Friedman and Eric J Alm. Inferring correlation networks from genomic survey data. PLoS computational biology, 8(9):e1002687, 2012.

9. Huaying Fang, Chengcheng Huang, Hongyu Zhao, and Minghua Deng. CCLasso: correlation inference for compositional data through lasso. Bioinformatics, 31(19):3172–3180, 2015.

10. Huaying Fang, Chengcheng Huang, Hongyu Zhao, and Minghua Deng. gcoda: conditional dependence network inference for compositional data. Journal of Computational Biology, 24(7):699–708, 2017.

11. Ksenia Guseva, Sean Darcy, Eva Simon, Lauren V Alteio, Alicia Montesinos-Navarro, and Christina Kaiser. From diversity to complexity: Microbial networks in soils. Soil Biology and Biochemistry, 169:108604, 2022.

12. Rosa Aghdam, Mojtaba Ganjali, Xiujun Zhang, and Changiz Eslahchi. CN: a consensus algorithm for inferring gene regulatory networks using the sorder algorithm and conditional mutual information test. Molecular BioSystems, 11(3):942–949, 2015.

13. Seyed Amir Malekpour, Maryam Shahdoust, Rosa Aghdam, and Mehdi Sadeghi. wplogicnet: logic gate and structure inference in gene regulatory networks. Bioinformatics, 39(2):btad072, 2023.

14. Gabriele Berg, Daria Rybakova, Doreen Fischer, Tomislav Cernava, Marie-Christine Champomier Vergé, Trevor Charles, Xiaoyulong Chen, Luca Cocolin, Kellye Eversole, Gema Herrero Corral, et al. Microbiome definition revisited: old concepts and new challenges. Microbiome, 8:1–22, 2020.

15. Alex D Washburne, James T Morton, Jon Sanders, Daniel McDonald, Qiyun Zhu, Angela M Oliverio, and Rob Knight. Methods for phylogenetic analysis of microbiome data. Nature microbiology, 3(6):652–661, 2018.

16. Catherine Lozupone, Manuel E Lladser, Dan Knights, Jesse Stombaugh, and Rob Knight. Unifrac: an effective distance metric for microbial community comparison. The ISME journal, 5(2):169–172, 2011.

17. Justin D Silverman, Alex D Washburne, Sayan Mukherjee, and Lawrence A David. A phylogenetic transform enhances analysis of compositional microbiota data. Elife, 6:e21887, 2017.

18. Chieh Lo and Radu Marculescu. Pglasso: Microbial community detection through phylogenetic graphical lasso. arXiv preprint 1807.08039, 2018.

19. Tony J Lam, Moses Stamboulian, Wontack Han, and Yuzhen Ye. Model-based and phylogenetically adjusted quantification of metabolic interaction between microbial species. PLoS computational biology, 16(10):e1007951, 2020.

20. Mehdi Layeghifard, David M Hwang, and David S Guttman. Disentangling interactions in the microbiome: a network perspective. Trends in microbiology, 25(3):217– 228, 2017.

21. Somnath Datta, Subharup Guha, et al. Statistical analysis of microbiome data. Springer, 2021.

22. Yang Ni, Veerabhadran Baladandayuthapani, Marina Vannucci, and Francesco C Stingo. Bayesian graphical models for modern biological applications. Statistical Methods & Applications, 31(2):197–225, 2022.

23. Jerome Friedman, Trevor Hastie, and Robert Tibshirani. Sparse inverse covariance estimation with the graphical lasso. Biostatistics, 9(3):432–441, 2008.

24. Nicolai Meinshausen and Peter Bühlmann. High-dimensional graphs and variable selection with the lasso. 2006.

25. Zachary D Kurtz, Christian L Müller, Emily R Miraldi, Dan R Littman, Martin J Blaser, and Richard A Bonneau. Sparse and compositionally robust inference of microbial ecological networks. PLoS computational biology, 11(5):e1004226, 2015.

26. Grace Yoon, Irina Gaynanova, and Christian L Müller. Microbial networks in spring-semi-parametric rank-based correlation and partial correlation estimation for quantitative microbiome data. Frontiers in genetics, 10:516, 2019.

27. Jian Xiao, Li Chen, Yue Yu, Xianyang Zhang, and Jun Chen. A phylogeny-regularized sparse regression model for predictive modeling of microbial community data. Frontiers in microbiology, 9:3112, 2018.

28. Antonio Gonzalez, Jose A Navas-Molina, Tomasz Kosciolek, Daniel McDonald, Yoshiki Vázquez-Baeza, Gail Ackermann, Jeff DeReus, Stefan Janssen, Austin D Swafford, Stephanie B Orchanian, et al. Qiita: rapid, web-enabled microbiome meta-analysis. Nature methods, 15(10):796–798, 2018.

29. Daphane Koller. Probabilistic graphical models: Principles and techniques, 2009.

30. Maryam Shahdoust, Rosa Aghdam, and Mehdi Sadeghi. Simmapnet: A bayesian framework for gene regulatory network inference using gene ontology similarities as external hint. bioRxiv, pages 2025–04, 2025.

31. Rosa Aghdam, Xudong Tang, Shan Shan, Richard Lankau, and Claudia Solís-Lemus. Human limits in machine learning: prediction of potato yield and disease using soil microbiome data. BMC bioinformatics, 25(1):366, 2024.

32. Donald T McKnight, Roger Huerlimann, Deborah S Bower, Lin Schwarzkopf, Ross A Alford, and Kyall R Zenger. Methods for normalizing microbiome data: an ecological perspective. Methods in Ecology and Evolution, 10(3):389– 400, 2019.

33. Li Chen, James Reeve, Lujun Zhang, Shengbing Huang, Xuefeng Wang, and Jun Chen. Gmpr: A robust normalization method for zero-inflated count data with application to microbiome sequencing data. PeerJ, 6:e4600, 2018.

34. Dieter Schlee. Numerical taxonomy. the principles and practice of numerical classification, 1975.

35. Emmanuel Paradis, Julien Claude, and Korbinian Strimmer. Ape: analyses of phylogenetics and evolution in r language. Bioinformatics, 20(2):289–290, 2004.

36. Carl Edward Rasmussen and Hannes Nickisch. Gaussian processes for machine learning (gpml) toolbox. The Journal of Machine Learning Research, 11:3011–3015, 2010.

37. Marta Karaliuė and Kestutis Dučinskas. Classification of gaussian spatio-temporal data with stationary separable covariances. Nonlinear Analysis: Modelling and Control, 26(2):363–374, 2021.

38. Sandra De Iaco, Donato Posa, Claudia Cappello, and Sabrina Maggio. On some characteristics of gaussian covariance functions. International Statistical Review, 89(1):36–53, 2021.

39. David K Duvenaud, Hannes Nickisch, and Carl Rasmussen. Additive gaussian processes. Advances in neural information processing systems, 24, 2011.

40. Eyal Krupka and Naftali Tishby. Incorporating prior knowledge on features into learning. In Artificial Intelligence and Statistics, pages 227–234. PMLR, 2007.

41. Andrew Gordon Wilson. Covariance kernels for fast automatic pattern discovery and extrapolation with Gaussian processes. PhD thesis, University of Cambridge Cambridge, UK, 2014.

42. Moreno Bevilacqua, Christian Caamaño-Carrillo, and Emilio Porcu. Unifying compactly supported and matérn covariance functions in spatial statistics. Journal of Multivariate Analysis, 189:104949, 2022.

43. Andrew Gelman, John B Carlin, Hal S Stern, and Donald B Rubin. Bayesian data analysis. Chapman and Hall/CRC, 1995.

44. S James Press. Applied multivariate analysis: using Bayesian and frequentist methods of inference. Courier Corporation, 2005.

45. Yi Zhang. Smart pca. In Twenty-First International Joint Conference on Artificial Intelligence, 2009.

46. Morris L Eaton and ML Eaton. Multivariate statistics: a vector space approach, volume 512. Wiley New York, 1983.

47. Chan-Fu Chen. Bayesian inference for a normal dispersion matrix and its application to stochastic multiple regression analysis. Journal of the Royal Statistical Society: Series B (Methodological), 41(2):235–248, 1979.

48. Adam J Rothman, Elizaveta Levina, and Ji Zhu. Generalized thresholding of large covariance matrices. Journal of the American Statistical Association, 104(485):177–186, 2009.

49. Rosa Aghdam and Claudia Solís-Lemus. CMiNet: An r package and user-friendly shiny app for constructing consensus microbiome networks. Methods in Ecology and Evolution, 17(1):52–66, 2026.

50. Qiyao Yang, Rosa Aghdam, Patricia Q Tran, Karthik Anantharaman, and Claudia Solis-Lemus. Unraveling keystone taxa: Interactions within microbial networks and environmental dynamics in lake mendota. bioRxiv, pages 2024–11, 2024.

51. Robert J Tibshirani and Bradley Efron. An introduction to the bootstrap. Monographs on statistics and applied probability, 57(1):1–436, 1993.

52. Duo Jiang, Courtney R Armour, Chenxiao Hu, Meng Mei, Chuan Tian, Thomas J Sharpton, and Yuan Jiang. Microbiome multi-omics network analysis: statistical considerations, limitations, and opportunities. Frontiers in genetics, 10:995, 2019.

53. Rosa Aghdam, Shan Shan, Richard Lankau, and Claudia Solís-Lemus. A hybrid framework for disease biomarker discovery in microbiome research combining bayesian networks, machine learning, and network-based methods. Biology Methods and Protocols, 11(1):bpaf089, 2026.

